# TWIST1 Modulates Cilia Length, Endocytic Vesicle Dynamics, and Cell-Cell Junctions during Neural Tube Morphogenesis

**DOI:** 10.1101/2025.10.03.680308

**Authors:** Derrick Thomas, Brittany M. Hufft-Martinez, Zarna Lalwani, Vi Pham, Mary Elmeniawi, An J. Tran, Jianming Xi, Irfan Saadi, Walid D. Fakhouri

## Abstract

**Background:** Endocytosis constitutes a fundamental cellular process governing development through coordinated regulation of plasma membrane remodeling and ciliogenesis, processes essential for cell shape changes and embryonic development. Although *Twist1* null embryos display complete cranial neural tube closure defects and conditional knockout in neuroectoderm disrupts cranial neural crest cell fate determination and delamination, the function of TWIST1 in neural tube morphogenesis remains unknown. We investigated the basis underlying neuroectodermal morphological abnormalities in TWIST1 mutant embryos, specifically the formation of ectopic lateral bending points and cellular disorganization, by examining TWIST1 function in cilia formation, adherens junction integrity, and endocytic vesicle dynamics.

**Results:** Immunofluorescence analysis revealed that cytosolic TWIST1 colocalizes with β-catenin and endocytic regulators LRP2 and RAB11B along the apical surface of cranial neuroectoderm. *Twist1* knockout resulted in reduced ciliary length and number. Quantitative PCR and Western blot analyses demonstrated upregulation of RAB11B and β-catenin at mRNA and protein levels in *Twist1* mutants. This molecular dysregulation coincided with increased accumulation of apical endocytic vesicles and altered expression profiles of endocytic component genes, ultimately modifying the apical neuroectodermal cell-cell junctions.

**Conclusion:** Our findings establish TWIST1 as a regulator of neuroectodermal morphology, demonstrating its ability to modulate ciliogenesis, endocytic vesicle dynamics, and cell-cell integrity.

## 1. INTRODUCTION

Craniofacial disorders are the second most common congenital birth defects worldwide, arising from disruptions in neural tube (NT) formation and cranial neural crest cell (CNCC) development. ^1^ The morphogenesis of NT and CNCC depends on complex and dynamic cellular and tissue rearrangements orchestrated by key morphogenetic processes, including cell proliferation, apical constriction, cell shape changes, cell intercalation, fate transition, and delamination, all critical for subsequent craniofacial development. ^2,3^ Most frontal craniofacial bone and cartilage structures are derived from CNCCs specified in the neural plate borders of the neuroectoderm. These specified cells undergo epithelial-to-mesenchymal transition (EMT), and delamination from neural fold edges to become migratory mesenchymal cells that reach the frontonasal process and pharyngeal arches where they differentiate into bone, cartilage, neurons, and smooth muscle. ^4-7^ While numerous contributory genes and signaling pathways regulating NT closure and CNCC formation have been identified, mechanistic understanding of the cellular and molecular modifications during cell fate transition and NT closure remains poorly understood.

TWIST1 is a basic helix-loop-helix transcription factor that serves as a master regulator of epithelial-to-mesenchymal transition (EMT) in various cancer cells, promoting metastasis and often contributing to chemoresistance. ^8-10^ During embryogenesis, *Twist1* regulates developmental genes involved in mesoderm differentiation, mesenchymal cell survival, osteogenic differentiation, and numerous cancer-associated pathways.^8^ Previous studies demonstrated that NT closure failure in *Twist1* null mice results from reduced proliferative capacity of adjacent mesodermal cells, which is critical for proper neural fold elevation and NT closure.^11^ However, recent studies have demonstrated that *Twist1* mRNA and protein are expressed in neuroectoderm during NT development, detected by scRNA-seq, and immunofluorescent staining,^12-14^ findings validated by quantitative in situ mRNA hybridization (Stereo-seq), suggesting a direct role in NT development and closure.^13^

Mutations in the human *TWIST1* gene can cause multiple common birth defects, including craniosynostosis, hypertelorism, ptosis, cleft palate, and limb abnormalities, while haploinsufficiency leads to Saethre-Chotzen syndrome.^15-21^ Notably, many of these disorders overlap with features observed in ciliopathies, genetic defects caused by dysfunction of the primary cilia, which present with craniofacial anomalies in approximately 30% of cases.^22,23^ Similarly, loss of function and knockout of *Twist1* in mice produce several overlapping phenotypes with human disorders, including cleft palate, craniosynostosis, and limb abnormalities.^12,18,24,25^ We previously demonstrated that *Twist1* cko in CNCCs and *Twist1* phospho-deficient mutant mouse embryos exhibit cranial neural tube clefts and craniofacial bone loss.^12^ We showed that TWIST1 is necessary for suppressing adherens junction proteins and epithelial markers during the epithelial-to-mesenchymal transition (EMT) of pre-migratory CNCCs as they emerge from neural plate borders. ^12^ These *Twist1* mutant mouse models confirm the critical function of TWIST1 in regulating NT formation.

Notably, TWIST1 expression overlaps with β-catenin at the apical surface of neuroectodermal cells and colocalizes with endocytic factors LRP2 and RAB11B in neuroectodermal cells.^14^ Furthermore, the cytoplasmic TWIST1 protein interacts with Tight Junction Protein 1 (TJP1) in cancer cells, and with δ-catenin in neuroectoderm and prostate cancer cells to stabilize cytoplasmic TWIST1 by ubiquitin modification.^26-28^ TWIST1 also functions as a mechano-mediator for matrix stiffness, which promotes nuclear translocation of TWIST1 by releasing it from the cytoplasmic binding partner Ras GTPase-activating protein-binding protein 1 (G3BP1).^26^ These findings proposed a new function of cytoplasmic TWIST1 through interactions with the β/δ-catenins and TJP1 to stabilize the protein and potentially modulate membrane-associated proteins before nuclear translocation. Therefore, the function of cytosolic TWIST1 and the importance of its interaction with β-δ-catenins require further molecular and cellular investigation during development and in cancer diseases.^12^

Endosomes are critical organelles formed by the cell membrane and Golgi apparatus, serving as vital regulators of signaling molecule internalization, translocation, ciliogenesis, and recycling of membrane-associated proteins.^29-32^ Among numerous genes associated with apical endosomal formation, *Lrp2* and *Rab11b*, represent well-studied examples in neuroectodermal cells. LRP2, a transmembrane receptor, plays a crucial role in NT closure. Endosomal proteins RAB11B and VANGL2 recycle the cell membrane-associated proteins within the apical surface of the neural folds to decrease apical surface area, enabling proper neuroectodermal closure.^14^ LRP2 and RAB11B are highly expressed along the apical surface with elevated expression at the dorsal midline and lateral edges.^*14*^ Endocytic trafficking proteins, such as RAB11B and RAB5 GTPase regulate important aspects of ciliary membrane delivery and turnover. Additionally, RAB proteins, including RAB34, have been implicated in modulating cilia length and function. Together, RAB GTPases play a critical role in coordinating vesicular trafficking, membrane delivery, and signaling events required for the assembly and maintenance of the primary cilium.^33-36^

This study investigates TWIST1’s impact on endosomal recycling proteins, ciliogenesis, and cell-cell junctions in neuroectodermal cells during NT closure and pre-migratory CNCC formation. We examined TWIST1 colocalization with endocytic vesicle markers, quantified mRNA and protein levels of membrane recycling proteins in *Twist1* null and conditional knockout mice, counted endocytic vesicles in *Twist1* mutant embryos, and performed quantitative analysis of cell-cell junctions to explain morphological and cellular changes in apical neuroectodermal cells. This study proposes a new function of TWIST1 in modulating primary ciliary length, endocytic vesicle dynamics, and cell-cell junctions in neuroectoderm during neural fold closure and CNCC formation.

## 2. RESULTS

### 2.1. Whole mount Images and histological analysis of neural tube abnormalities

A whole mount image of *Twist1* null embryo exhibits complete neural tube closure defects with multiple ectopic bending points, and *Twist1* cko embryo presents neural tube developmental defects and improper closure compared to wild-type littermates at E10.5 (Figure 1A-D). To further analyze morphological changes, we performed H&E staining on sections of mouse neural folds to identify morphological changes in *Twist1* null and cko embryos. In control embryos (WT), the neural folds showed proper bending at the ventral midline and at the lateral dorsal regions important for NT closure (Figure 1E-E’, G-G’), while the *Twist1* null embryos exhibited multiple folding and elevation at the middle-lateral regions without the formation of proper bending points leading to complete failure of NT closure (Figure 1F-F’, H-H’). Similarly, control embryos showed proper folding and bending at the lateral dorsal regions (Figure 1II’, K-K’-). The *Twist1* cko embryos showed narrow ventral midline bending and multiple lateral dorsal bending points. Also, the expansion of the neural folds varied from the dorsal to the midline regions (Figure 1J-J’, L-L’).

**FIGURE 1.**
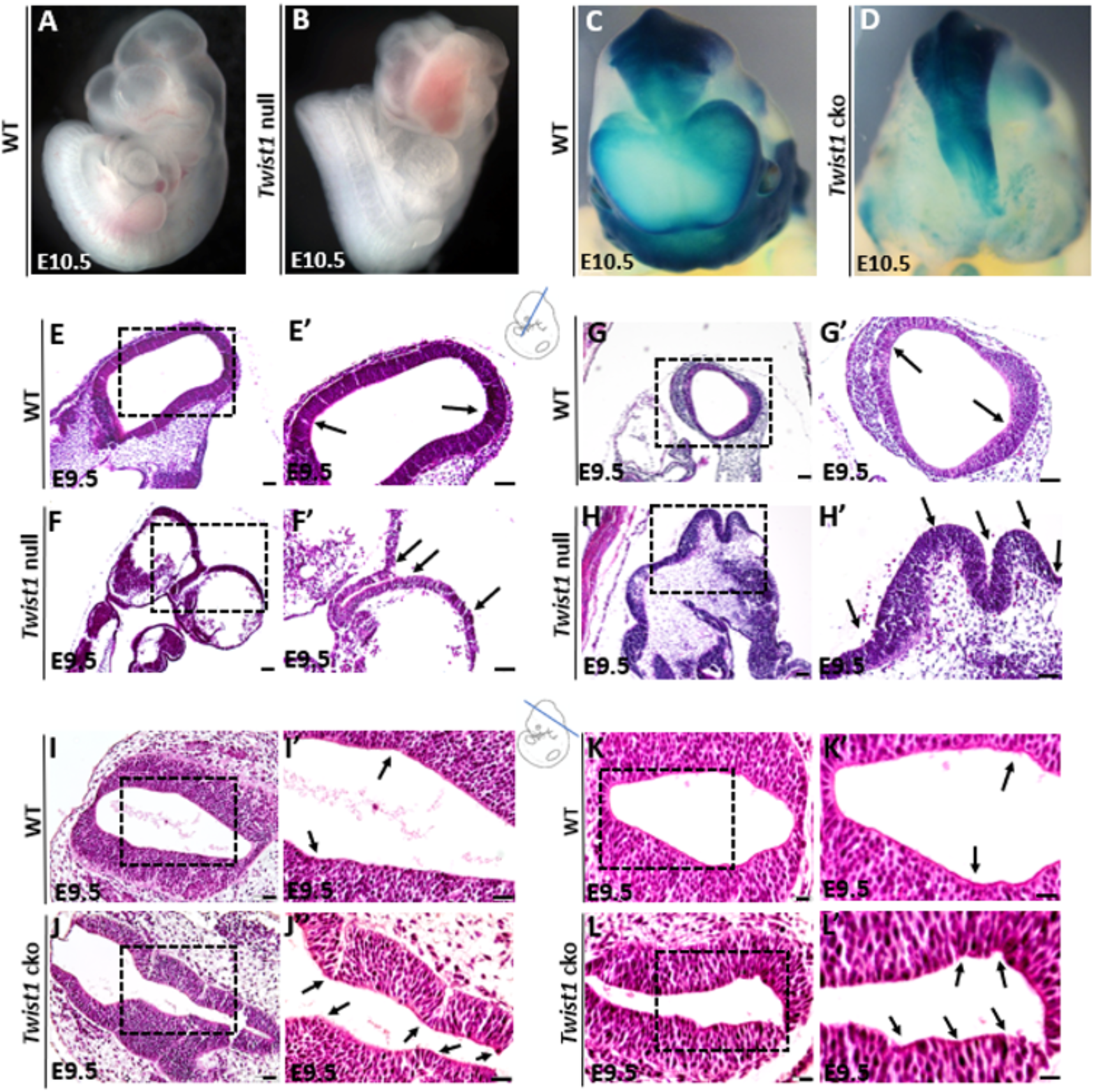
Whole mount Embryos and Histological analysis of neural fold development in *Twist1* mutant embryos. **(A):** Wildtype embryo at E10.5 showing a normal development and closure of neural tube. **(B):** *Twist1* null embryo exhibiting complete neural tube closure defects at E10.5. **(C):** Wildtype embryo at E10.5 showing a normal development and closure of neural tube (CNCC TM1). **(D):** *Twist1* cko embryo at E10.5 presents neural tube developmental defects and improper closure. **(E, E’):** H & E staining of E9.5 wildtype mouse embryo showing the morphology and closure of midbrain NT with organized bending points (black arrows) of neuroectodermal brain cortex. **(F, F’):** H & E staining of E9.5 *Twist1* null mouse exhibiting abnormal morphology and disorganized folding of the neuroectodermal folds. **(G, G’):** H & E staining of E9.5 wildtype mouse showing the morphology and closure of the hindbrain NT. **(H, H’):** H & E staining of E9.5 *Twist1* null mouse showing the severe abnormal morphology and disorganized folding of the mid- and hindbrain of neuroectodermal folds. **(I, I’):** H & E staining of E9.5 wildtype mouse showing proper morphology and closure of the hindbrain NT. **(J, J’):** H & E staining of E9.5 *Twist1* cko mouse showing the abnormal morphology of the neural folds with multiple bending points and expansion of neuroectoderm (black arrows). **(K, K’):** H & E staining of E9.5 wildtype mouse showing proper morphology and closure of the midbrain NT. **(L, L’):** H & E staining of E9.5 *Twist1* cko mouse showing the abnormal morphology of the neural folds with multiple bending points of neuroectoderm (black arrows).

### 2.2. TWIST1 expression in neural folds

Dual immunofluorescence (IF) staining of TWIST1/RAB11B, TWIST1/LRP2, RAB11B/β-catenin and LRP2/β-catenin demonstrated that TWIST1 is expressed at the apical region of wild-type E9.5 and E10.5 mouse embryos’ dorsal neuroectodermal cells, and its expression overlaps with the apical endocytic markers RAB11B and LRP2 as indicated by the yellow fluorescent color (Figure 2A-D”). We also detected a strong nuclear expression of TWIST1 in the delaminated CNCCs along the neural folds and near the bending sites (Figure 2A, B, D). The expression of adherens junction protein β-catenin also overlaps at the dorsal lateral region with the endocytic vesicle marker LRP2 (Figure 2E-E”), and RAB11B (Figure 2F-F”) at the apical side of the neuroectodermal cells.

**FIGURE 2.**
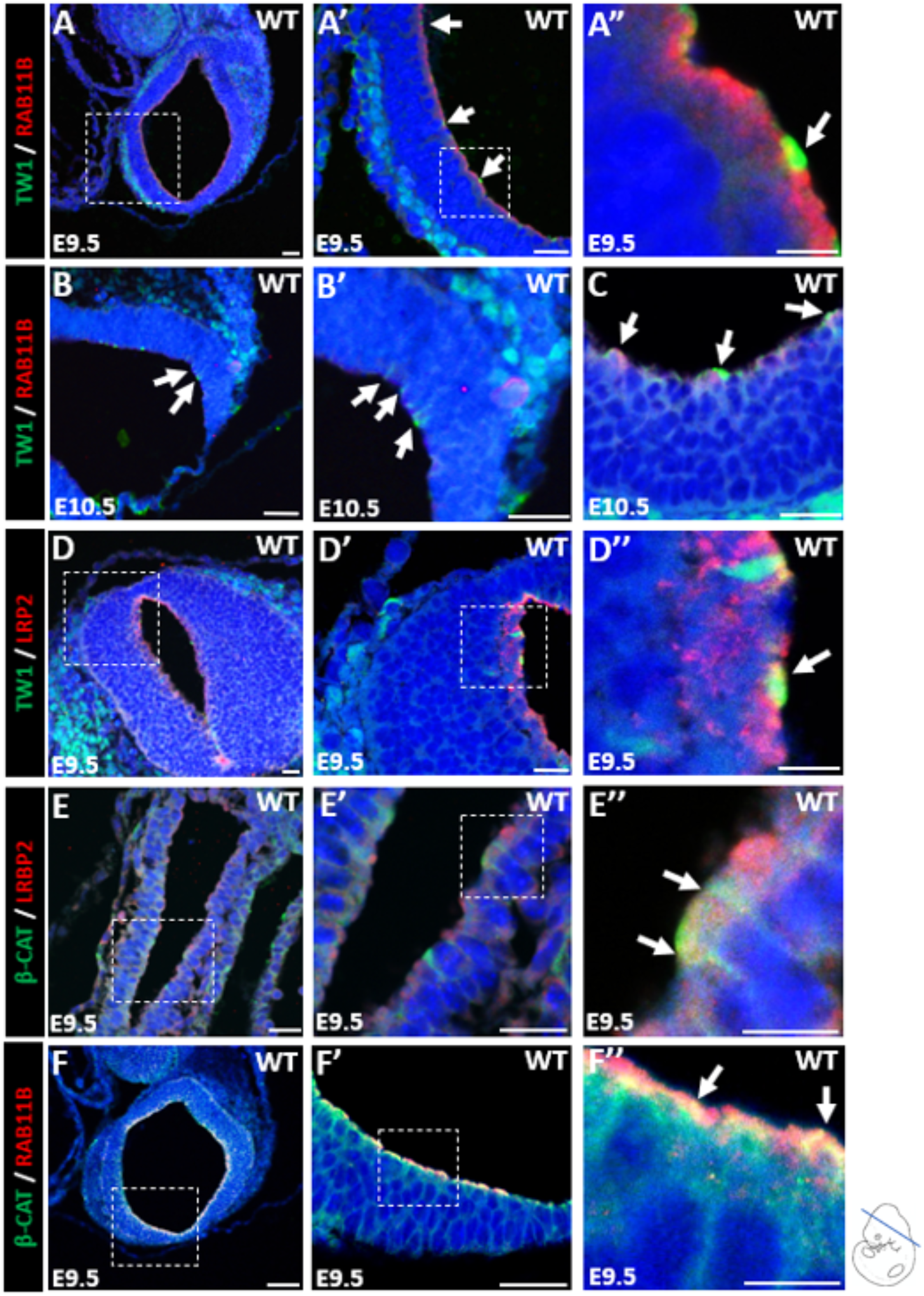
Expression pattern of TWIST1, endocytic vesicle markers, and adherens junction protein. **(A-A’’):** Immunostaining of E9.5 wildtype mouse embryo showing TWIST1 protein expression in green at apical surface of neuroectodermal cells, and its expression overlaps with endocytic vesicle marker, RAB11B, in yellow (white arrow). TWIST1 nuclear expression was detected in the delaminated cranial neural crest cells along the NT. **(B-B’, C):** Immunostaining of E10.5 wildtype mouse embryo showing TWIST1 protein expression in green at apical surface of neuroectodermal cells, and its expression overlaps with endocytic vesicle marker, RAB11B, in yellow (white arrow). TWIST1 nuclear expression was detected in the delaminated cranial neural crest cells along the NT and near bending sites.**(D-D’’):** Immunostaining of E9.5 wildtype mouse embryo showing TWIST1 protein expression in green at apical surface of neuroectodermal cells, and its expression overlaps with endocytic vesicle marker, LRP2, in yellow (white arrow). **(E-E’’):** Immunostaining of E9.5 wildtype mouse embryo showing the expression of β-catenin in green at apical-lateral side of neuroectodermal cells, and its expression overlaps with endocytic vesicle marker, LRP2, in yellow (white arrow). **(F-F’’):** Immunostaining of E9.5 wildtype mouse embryo showing the expression of β-catenin in green at apical-lateral side of neuroectodermal cells, and its expression overlaps with endocytic vesicle marker, RAB11B, in yellow (white arrow).

### 2.3. Shortened cilia in *Twist1*^*-/-*^ neuroectoderm

We investigated primary cilia length using ARL13B, ciliary membrane marker, to determine TWIST1 impact on ciliary length utilizing control and *Twist1*^*-/-*^ tissue from E9.0 and 9.5 embryos (Figure 3A-B’). We measured primary cilia length and found a significant reduction in *Twist1*^*-/-*^ cilia length compared to control (Figure 3C). We next looked at the ciliation in *Twist1*^*-/-*^ tissue from E9.0 and E9.5 embryos, and observed a significant decrease in the number of ciliated cells compared to wild-type littermates (Figure 3D). This decrease in *Twist1*^*-/-*^ cilia length and percent ciliated cells suggests TWIST1 is necessary for normal ciliogenesis in the neuroectoderm.

**FIGURE 3.**
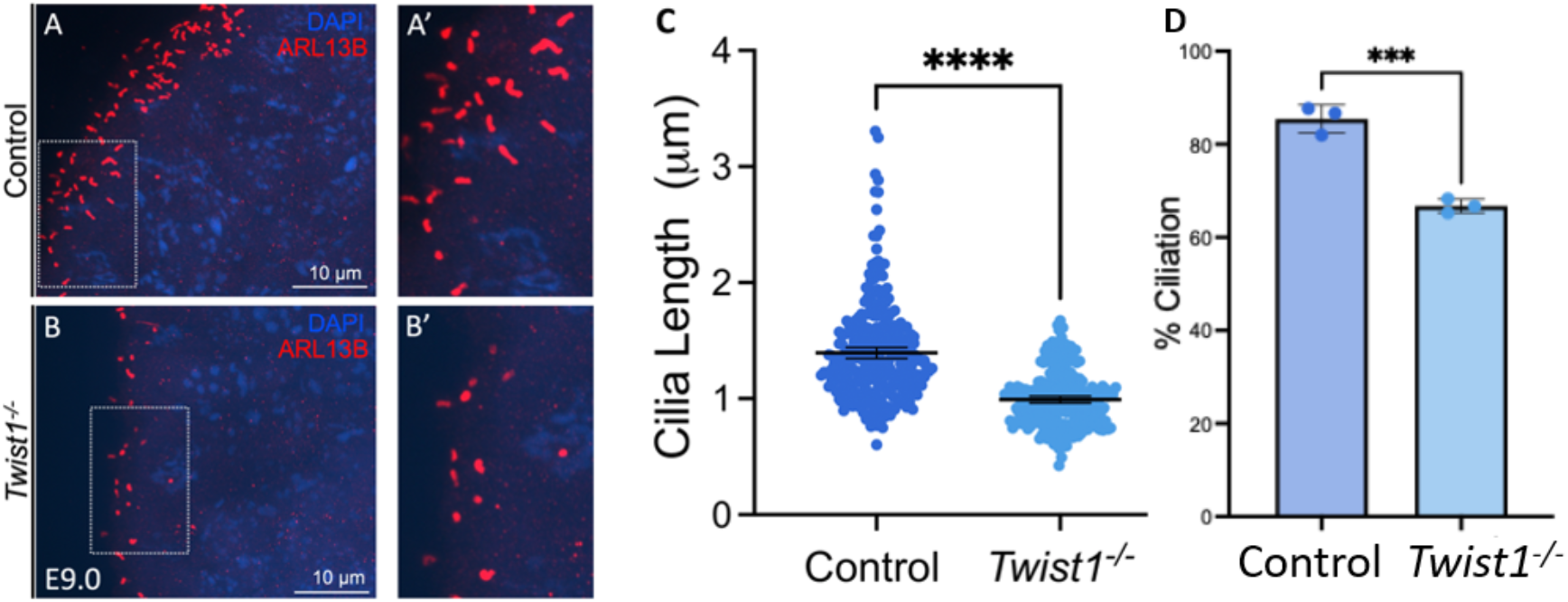
Primary cilia length is decreased in *Twist1*^*-/-*^ neuroectoderm. Immunostaining with ARL13B of control (A) and *Twist1*^*-/-*^ (B) mouse neuroectoderm from E9.0 embryos. **(A’-B’):** Magnification of boxed regions highlighting primary cilia. **(C):** Quantification of cilia length (μm) from E9.0 - 9.5 embryonic neurectoderms stained with ARL13B. Approximately 50 primary cilia per embryo were measured from 6 control (n=300) and 5 *Twist1*^*-/-*^ (n=239) samples. (**D**): Quantification of percent ciliation from E9.0-9.5 embryonic neuroectoderms stained with ARL13B and DAPI. For percent ciliation quantification, 3 control (n=530) and *Twist1*^*-/-*^ (n=734) samples were measured.

### 2.4. Quantification of mRNA and protein of endocytic and adhesion junction markers in *Twist1* null and cko

We observed no significant changes in LRP2 mRNA transcripts in *Twist1*^*-/-*^ and *Twist1*^*cko/-*^ compared to wild type at E8.5, E9.0, E9.5, and E10, whereas an almost two-fold increase in expression of RAB11B and β-catenin was detected at E8.5, E9.0, and E9.5 embryonic stages (Figure 4A-I). However, no significant changes in expression levels of RAB11B or β-catenin were observed for *Twist1*^*cko/-*^ at E10.0 (Figure 4J-L).

**FIGURE 4.**
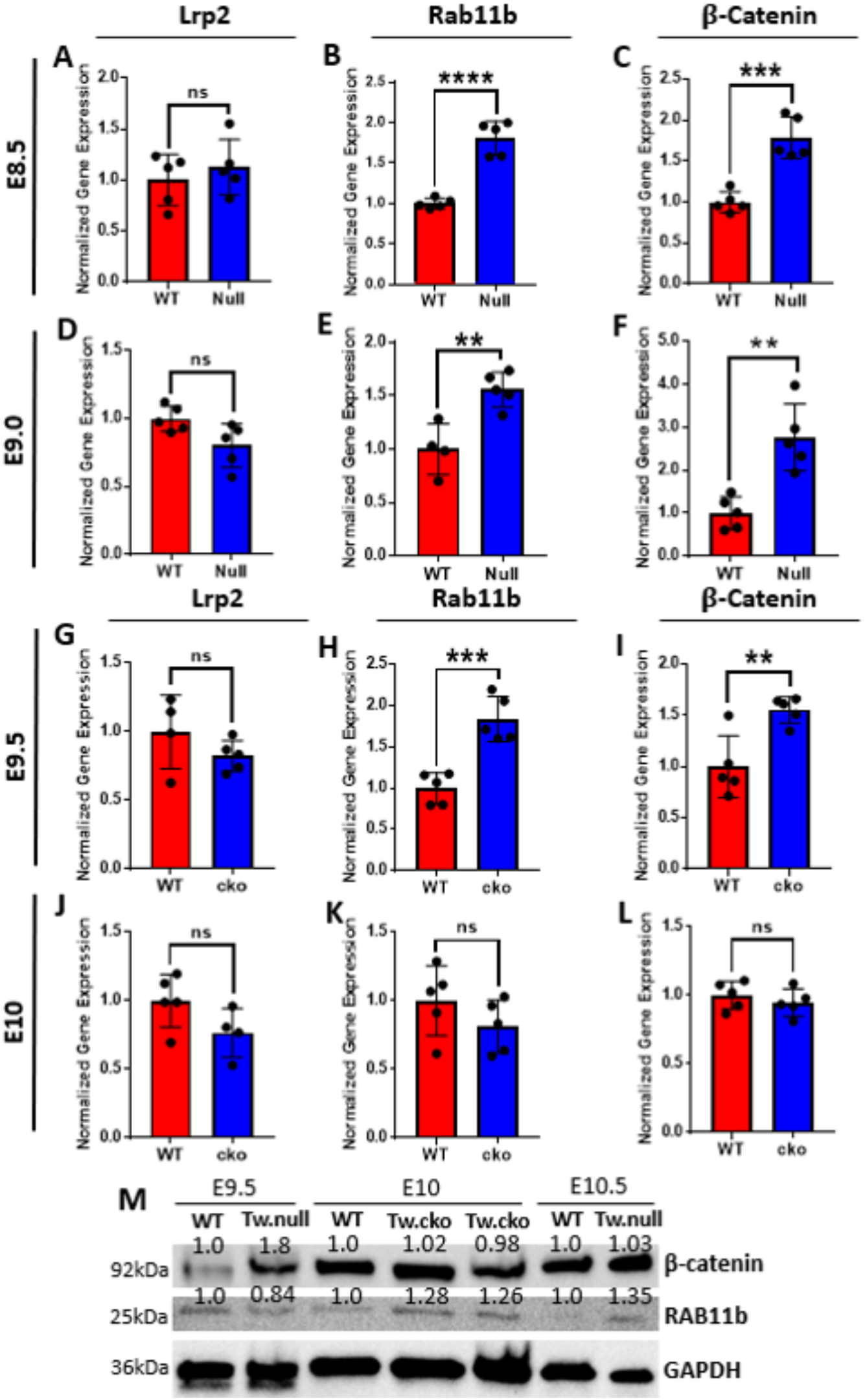
Quantitative analysis of endocytic vesicle markers and β-catenin in *Twist1* mutant embryos. **(A-F):** mRNA expression of *Lrp2, Rab11b*, and β*-catenin* in hindbrain of *Twist1* null embryos at E8.5 and E9. The Expression of *Rab11b* and β*-catenin* significantly increased while no change in *Lrp2* was detected. **(G-L):** mRNA expression of *Lrp2, Rab11b*, and β*-catenin* in hindbrain of *Twist1* cko embryos at E9.5 and E10. The Expression of *Rab11b* and β*-catenin* significantly increased at E9.5 while no significant change was detected in the expression of *Lrp2, Rab11b*, and β*-catenin* at E10 between mutant and wildtype mouse embryos. For statistical analysis, the * represents a p-value < 0.05, ** represents a p-value < 0.01, *** represents a p-value < 0.001, and ****represents a p-value < 0.0001. **(M):** Protein level of β-catenin and RAB11B was quantified in *Twist1* null and cko compared to corresponding WT embryos and the protein level was normalized to the housekeeping gene GAPDH. A slight increase in the protein level of both *Twist1* null and cko was detected compared to WT for β-catenin and RAB11B.

To evaluate the changes at the protein level, a western blot was performed for total proteins extracted from *Twist1*^*-/-*^ and *Twist1*^*cko/-*^ samples and compared to the corresponding wild type littermates. At E9.5, total β-catenin protein was increased in *Twist1*^*-/-*^ when compared to WT. A similar trend was observed at embryonic stage E10 and E10.5 in *Twist1*^*-/-*^ and *Twist1*^*cko/-*^, the level of β-catenin and RAB11B was elevated compared to WT. This shows that both β-catenin and RAB11B are expressed in earlier embryonic time points and their expression elevated in *Twist1* mutant embryos. We used the GAPDH as a loading control and for normalization (Figure 4M).

### 2.5. Immunofluorescent staining of endocytic markers

To determine the expression pattern of endocytic vesicle markers, dual IF staining for LRP2 and β-catenin as well as RAB11b and β-catenin was performed on wild type, *Twist1* null, and cko embryos at E9.5 and E10.5 (Figure 5 and S1). The IF images show that RAB11B and LRP2 are highly expressed in apical cells in both cko and null neuroectoderm when compared to corresponding WT tissues (Figure 5A-H’). However, the cells beneath the dorsal apical cells didn’t show robust expression of LRP2 and RAB11B, particularly in wild type (Figure 5A’, C’, E’, G’). The increase in staining in mutant embryos suggests an elevation in the number of apical endosomes in the absence of *Twist1*. To investigate the level of change further, we counted the number of apical endocytic vesicles as a measure of quantitative analysis for endosome dynamic.

**FIGURE 5.**
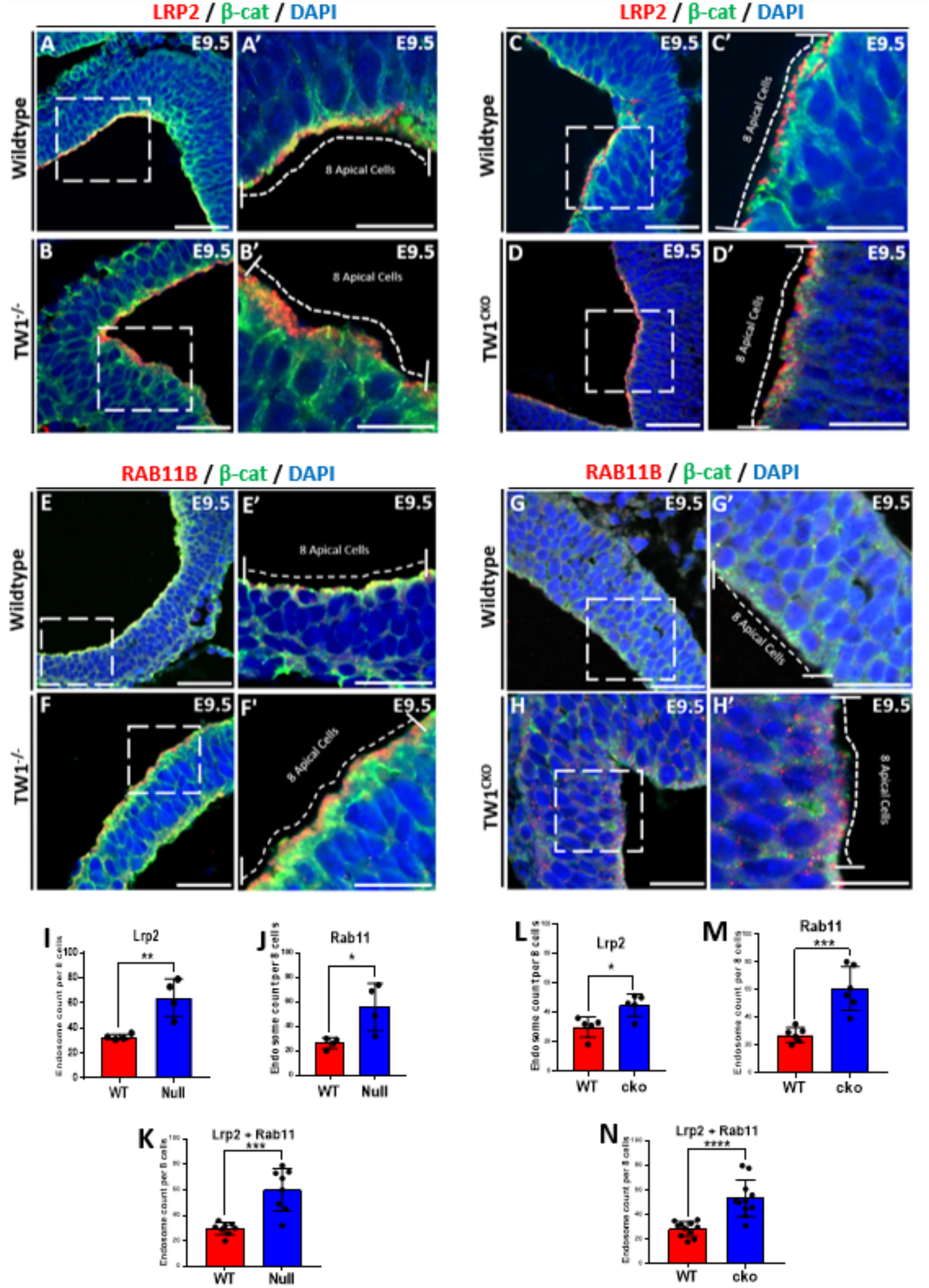
Expression pattern and quantification of endocytic vesicles in neuroectodermal cells. **(A-B’):** IF staining of LRP2 and β-catenin in E9.5 *Twist1* null and WT littermates. The endocytic vesicles were quantified in 8 apical cells where β-catenin was used as cell membrane marker to identify cell peripheries. **(C-D):** IF staining of LRP2 and β-catenin in E9.5 *Twist1* cko and WT littermates. **(E-F’):** IF staining of RAB11B and β-catenin in E9.5 *Twist1* null and WT littermates. **(G-H’):** IF staining of RAB11B and β-catenin in E9.5 *Twist1* cko and WT littermates. **(I-K):** The number of endocytic vesicles was increased in *Twist1* null using LRP2 and RAB11B and also in the combined quantification of both markers. **(L-N):** The number of endocytic vesicles was increased in *Twist1* cko using LRP2 and RAB11B and also in the combined quantification of both markers. For statistical analysis, the * represents a p-value < 0.05, ** represents a p-value < 0.01, *** represents a p-value < 0.001, and ****represents a p-value < 0.0001.

### 2.6. Impact of *Twist1* on the Endocytic Vesicle Dynamic

To analyze changes in the number of endocytic vesicles, we counted the number of endocytic vesicles in 8 apical neuroectodermal cells of IF images using the two markers, LRP2 and RAB11B. We used β-catenin as a cell-membrane associated marker to determine the cell peripheries. IF images of *Twist1* null and cko embryos at time points E9.5 and E10.5 depict a considerable increase in endosomal markers LRP2 (Figure 5A-D’ and S1) and RAB11B compared to WT (Figure 5E-H’). Most of the endocytic vesicles were found at the apical surface of the dorsal neuroectodermal cells in the WT, while for the null and cko, they were moderately shifted more towards the apical lateral regions. When we counted the apical endosomes for LRP2 in 8 apical cells, we found that the number of endosomes increased around two-fold in *Twist1* null and cko compared to wild-type littermates of sections of the hindbrain (Figure 5I, L). A similar trend was seen in both *Twist1* null and cko for RAB11B (Figure 5J, M). When combined the quantification for both biomarkers, the LRP2 and RAB11B, *Twist1* null and cko showed a significant increase in apical endosomes which is consistent with the staining in individual IF images (Figure 5K, N). Our data demonstrates that loss of TWIST1 significantly increase the number of endocytic vesicles at the apical surface of neuroectoderm during CNCC formation.

### 2.7. mRNA expression of other endocytic regulator proteins

Apart from *Lrp2* and *Rab11b*, other endosomal genes involved in endosomal formation and recycling were also evaluated in *Twist1* null at E9.0 and *Twist1* cko at E9.5. We extracted RNA from hindbrain and first pharyngeal arch of WT and mutant embryos for quantification. We used three biological replicates for either WT and mutant and five technical replicants for each genotype for the RT-qPCR assay. In *Twist1* null embryos, the expression of Dynamin1 (*Dnm1) and* Ubiquitin-specific peptidase 2 *(Usp2)* was significantly higher compared to WT littermates at E9.0 (Figure 6A, G). No statistically significant difference was found in the expression of Actin-Related Proteins 2 and 3 (*Arp2* and *Arp3)* and Tight junction protein 1 (*Tjp1)* between *Twist1* null embryos and WT (Figure 6B-D). However, we observed a decrease in the expression of VANGL Planar Cell Polarity Protein 2 *(Vangl2)* and Clathrin (*Cltc)* in *Twist1* null compared to WT samples (Figure 6E, F). When compared to WT, the level of *Dnm1* expression increased more than three-fold in *Twist1* cko (Figure 6H). *Arp2* and *Arp3* showed a significant increase, whereas *Tjp1* expression showed a nearly 50% reduction in *Twist1* cko (Figure 6I-K). However, the expression levels of *Vangl2, Cltc*, and *Usp2* remain similar in WT and *Twist1* cko samples (Figure 6L-N).

**FIGURE 6.**
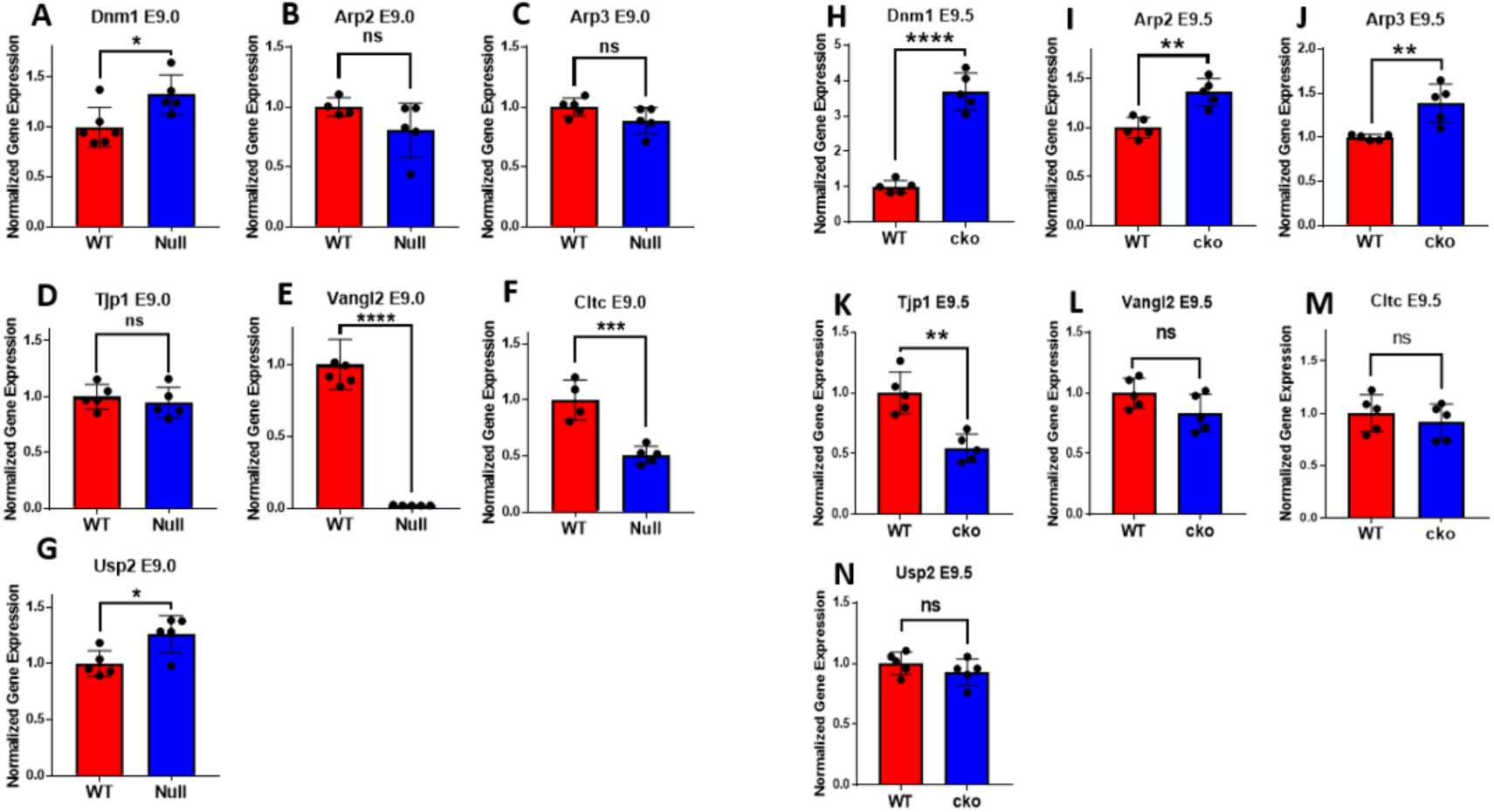
mRNA quantification of endocytic vesicle markers and associated genes. **(A-G):** A significant increase in the expression level of *Dnm1* and *USP2* was detected while the expression of *Vangl2* and *Cltc* was significantly decreased in E9.0 *Twist1* null compared to WT littermates. There was no significant change detected in remaining tested genes between mutant and WT embryos. **(H-N):** A significant increase in the expression level of *Dnm1, Arp2*, and *Arp3* was detected while the expression of *Tjp1* was significantly decreased in E9.0 *Twist1* cko compared to WT littermates. There was no significant change detected in remaining tested genes between mutant and WT embryos. For statistical analysis, the * represents a p-value < 0.05, ** represents a p-value < 0.01, *** represents a p-value < 0.001, and ****represents a p-value < 0.0001.

### 2.8. Quantification of the cell-cell interface in Twist1 null and cko embryos

To determine the alternations in cell morphology of neuroectoderm during NT development, IF images for β-catenin at multiple time points were analyzed using the Junction Mapper software. Cell-cell interfaces for neuroectodermal cells near the apical side of the neural folds were analyzed based on interface contour, interface linearity index, interface area, junction marker intensity, and junction marker intensity per area. Images at the edges of the neural folds were chosen for *Twist1*^*-/-*^ with additional images included in the Supporting Materials (Figure 7A-D and S2, 3, 4). A significant decrease in interface contour and interface area was observed in *Twist1*^*-/-*^ compared to the wild type (Figure 7E, F). However, there was a significant increase in the interface linearity index between *Twist1*^*-/-*^ and wild type, and no significance in JM1 intensity and JM1 intensity per area (Figure 7G-I). Images at the edges of the neural folds were also selected for *Twist1*^*cko/-*^ with the corresponding WT (Figure 7J-M). A significant decrease in interface contour and interface area was observed for *Twist1*^*cko/-*^ compared to the wild type (Figure 7N, O). There was no significant difference in the interface linearity index and JM1 intensity (Figure 7P, Q), while there was a significant increase in JM1 intensity per area of *Twist1*^*cko/-*^ compared to wild type (Figure 7R).

**FIGURE 7.**
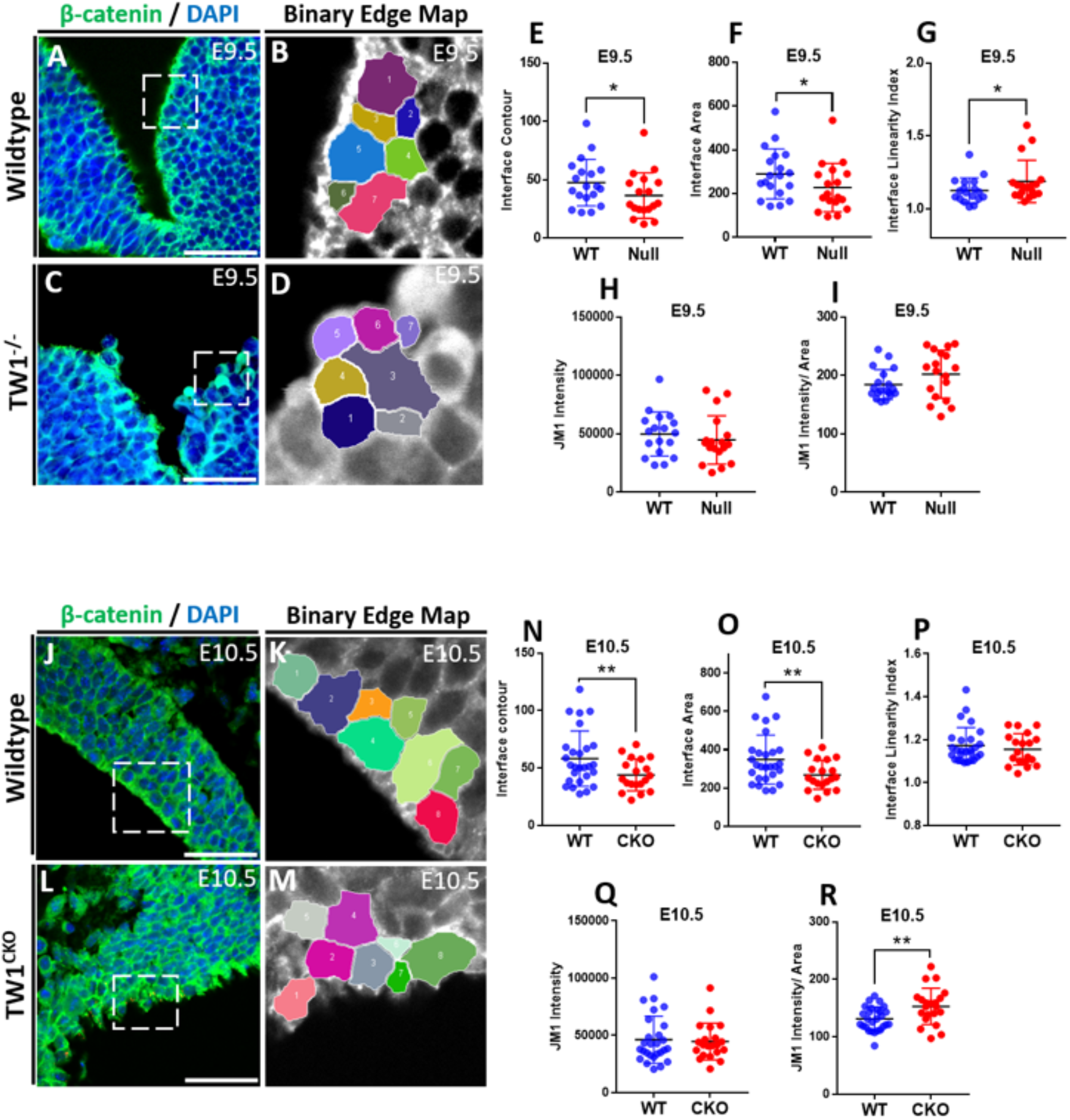
Quantification analysis of cell-cell junction of neuroectodermal cells of Twist1 mutants and WT embryos. **(A):** IF staining of E9.5 WT mouse embryo using β-catenin as cell membrane marker. **(B):** Binary edge map of E9.5 WT mouse embryo of the cells used for quantification analysis. **(C):** IF staining of E9.5 *Twist1* null mouse embryo using β-catenin as cell membrane marker. **(D):** Binary edge map of E9.5 *Twist1* null mouse embryo of the cells used for quantification analysis. **(E-I):** Quantification analysis shows a decrease in the interface contour and interface area while there is an increase in the interface linearity index in *Twist1* null compared to WT. **(H, I):** There is no significant change detected in JM1 intensity and JM1 intensity / area between the mutant and WT embryos. **(J):** IF staining of E10.5 WT mouse embryo using β-catenin as cell membrane marker. **(K):** Binary edge map of E10.5 WT mouse embryo of the cells used for quantification analysis. **(L):** IF staining of E10.5 *Twist1* cko mouse embryo using β-catenin as cell membrane marker. **(M):** Binary edge map of E10.5 *Twist1* cko mouse embryo of the cells used for quantification analysis. **(N-R):** Quantification analysis shows a significant decrease in the interface contour and interface area while there is an increase in JM1 intensity / area in *Twist1* cko compared to WT. There is no significant change detected in interface linearity index and JM1 intensity between the mutant and WT embryos. Six biological replicates were used for IF staining and around 200 technical replicates were used for the cell-cell contact data analysis. For statistical analysis, the * represents a p-value < 0.05, ** represents a p-value < 0.01, *** represents a p-value < 0.001, and ****represents a p-value < 0.0001.

## 3. DISCUSSION

Embryonic development is orchestrated through complex molecular and cellular processes governed by intricate interactions between genetic determinants, environmental influences, and coordinated signaling networks that direct cell lineage specification and differentiation^37^. The multifaceted regulatory interactions within embryonic cells, combined with our incomplete mechanistic understanding, present significant challenges in elucidating the causative factors of craniofacial abnormalities, which represent the second most prevalent category of congenital birth defects.^16-21,24^ Identifying the regulatory pathways and associated cellular changes that drive tissue morphogenesis is therefore crucial for addressing fundamental biological concepts in developmental processes and the molecular basis of craniofacial disorders.

During neurulation, mesodermal signals induce the overlying dorsal ectoderm to differentiate into neural and non-neural ectoderm, establishing specified cell populations at the neural plate borders through tightly regulated gene expression programs essential for cell fate determination^3,37^. Subsequently, the neural tube develops into the entire central and peripheral nervous system^38,39^. This study aimed to elucidate TWIST1-dependent molecular and cellular alterations in neuroectoderm and determine why *Twist1* deficiency causes cellular disorganization and multiple ectopic lateral bending points along the dorsal neural folds leading to neural tube closure defects and brain abnormalities. We investigated whether TWIST1 colocalizes with adherens junction and endosomal proteins in neuroectodermal cells and whether *Twist1* loss alters primary ciliary length, endosomal marker expression, β-catenin levels, endocytic vesicle dynamics, and cell junction organization in neuroectodermal cells at dorsolateral regions. Our goal was to uncover unknown molecular and cellular alterations associated with neural tube defects in *Twist1* mutant mouse models. While our current data do not establish the precise mechanism by which TWIST1 impacts endocytic vesicle dynamics and ciliogenesis, they further our previous findings that demonstrated that TWIST1 protein interacts with cytosolic β- and δ-catenins as part of the adherens junction protein complex which affects cell shape changes and membrane activities. ^12^.

Recent transcriptomic analyses and quantitative in situ hybridization studies have demonstrated TWIST1 expression in neural plate borders and neural folds,^13,40^ providing compelling evidence for its direct functional role in neuroectodermal cellular development. Our immunostaining validated moderate TWIST1 expression in neuroectodermal cells during NT development. *Twist1* null and conditional knockout mouse models exhibit failed cranial neural tube closure, formation of multiple ectopic bending points, and disrupted cranial neural crest cell fate transition and survival, resulting in significant craniofacial bone deficiencies^12^. To determine TWIST1’s impact on primary ciliogenesis, we examined cilia length and number using the ciliary membrane marker ARL13B. Our data revealed significantly reduced ciliary length at the dorsal *Twist1* null neuroectoderm compared to control littermates at E9-9.5. The number of ciliated cells also appears to be reduced in mutant neuroectodermal cells. Alterations in endocytic vesicle protein expression and β-catenin levels, along with increased endocytic vesicle abundance, could potentially explain the shortened cilia in mutant embryos. These findings suggest that TWIST1 is involved in adherens junction integrity and endocytic vesicle dynamics, which could explain the changes observed in cell-cell junction and intercellular spatial organization. These cellular processes facilitate apical constriction, membrane-associated protein recycling, and cell fate transitions during neurulation^12,41^.

In TWIST1’s absence, neuroectodermal disorganization accompanied by multiple bending points and shorter cilia may help explain neural tube closure defects in *Twist1* null embryos and exencephaly in *Twist1* conditional knockouts^12^. The altered expression of genes involved in endocytic vesicle function and elevated endocytic vesicle numbers are consistent with observed increase in LRP2 and RAB11B proteins in *Twist1* null and conditional knockout hindbrain embryos. Our previous and current findings demonstrate the TWIST1’s integral role in neural tube closure and cell fate transition of pre-migratory CNCC^12^. We demonstrate that TWIST1 activity directly modulates neural fold bending and cell-cell junctions in regions where pre-migratory cranial neural crest cells delaminate from neural fold edges. The significance of these findings is underscored by the current absence of direct evidence linking TWIST1 dysfunction to cranial neural tube closure defects in human cases, despite the fact that the etiological basis for most human neural tube defects, including anencephaly and exencephaly, remains largely undefined^42^. Based on our findings, we suggest a link between *TWIST1* locus and downstream target genes involved in neural tube development. Interestingly, recent examination of *TWIST1* regulatory elements led to the identification of mutations linked to Auriculocondylar syndrome ^43^ and limb malformations^44^. These investigations may reveal previously unrecognized connections between TWIST1 dysfunction and birth defects associated with neural tube disorders, potentially opening avenues for diagnostic and therapeutic interventions.

## 4. EXPERIMENTAL PROCEDURE

### 4.1. Mouse Strains

All mice used in this study were generated using C57BL/6J mouse genetic background. *EIIA-Cre* mouse line was crossed to the B6;129S7-*Twist1*^*fl/fl*^ mice to obtain the *Twist1* heterogeneous mice, as previously described.^12^ *Twist1* cko in neuroectoderm was generated using the *Wnt1-Cre2* deleter strain. The number of experimental animals was calculated based on the power analysis. The animal work was approved by the CLAMC committee at UTHealth Houston under the Approved Animal protocol number AWC-22-0060.

### 4.2. Histological and Immunofluorescent Staining

Embryonic tissues from control, *Twist1*^*-/-*^, and *Twist1*^*cko/-*^ mice at E9-9.5 were embedded in paraffin, sectioned, and stained with hematoxylin and eosin (H&E) for histological analysis. For immunofluorescence staining, homozygous *Twist1 null* and cko mutant embryos at E9-9.5 were identified via phenotyping of the neural tubes, either open or malformed neural folds. Littermate embryos with phenotypically normal neural tubes were used as control. The wild-type *Twist1*^*+/+*^, *Twist1*^*cko/-*^, and *Twist1*^*-/-*^ embryos at later timepoints were genotyped via PCR as described previously^45^. Embryos were collected for detecting the expression pattern and level of endocytic markers and adherens junction proteins. Expression of LRP2, RAB11b, and β-catenin was performed on head sections of paraffin-embedded mouse embryos. The immunofluorescent staining protocol was followed as previously described.^45^ The primary antibodies used for the staining are monoclonal TWIST1 (1:150, Abcam, MA), mouse monoclonal anti-β-catenin (1:200, Santa Cruz), rabbit monoclonal anti-RAB11B (1:130, ABclonal), and rabbit monoclonal anti-LRP2 (1:150, ABclonal). The sections were incubated for 3 hours with secondary antibodies conjugated to Alexa fluorophores 488 goat anti-rabbit (1:150) and 555 goat anti-mouse (1:150). The sections were counterstained for nuclei with water-diluted DAPI. The mounted sections were imaged using a Nikon C2 Confocal microscope.

### 4.3. Cilia Staining and Measurements

E9.0-E9.5 embryo sections were used for cilia staining. Antigen retrieval was performed by heating slides in sodium citrate buffer (10 mM sodium citrate, 0.05% Tween 20, pH 6.0) at 96°C for 10 min. Slides were washed in PBS, permeabilized using 0.5% Triton X-100 in 1× PBS for 30 min, washed in PBS again, then blocked in 10% NGS. Primary and secondary antibodies were incubated following the same method as described above. Primary antibody for ARL13B (1:300, Proteintech, 17711-1-AP) and secondary antibody of Goat anti-Rabbit IgG (H + L) Alexa 594 (1:500 tissue Invitrogen; A-11037) were used. DAPI (5 μM) was used to mark nuclei. Images were acquired using the Nikon CSU-W1 Spinning-disk confocal 60x objective and an intermediate 4x magnification changer with SoRa module. A z-stack was captured with slices taken every 0.2μm ensuring the entire tissue section was captured. A max intensity projection (MIP) was created for measurement. Cilia lengths were measured using ImageJ software and a segmented line was used to trace the cilium^46,47^.The length in pixels was converted to microns using the pixel conversion factor for the objective used. MIP image was used to count the % ciliation in the neuroectoderm.

### 4.4. RT-qPCR

The mRNA expression of *Lrp2, Rab11b*, and *β-catenin* in the mid-and hindbrain regions of mouse embryos, was measured in three pooled biological and five technical replicates at E8.5 and E9.0 for *Twist1* null embryos, and E9.5 and E10 for *Twist1* cko compared to wildtype littermates. We also quantified the expression of genes involved in endocytosis, including *Vangl2, Cltc, Dnm1, Tjp1, Usp2, Arp2*, and *Arp3*.

### 4.5. Western Blot Assay

Midbrain and hindbrain tissues were dissected from embryos at E9.5, E10, and E10.5 and used to extract total protein. Tissues were directly frozen and thawed twice after grinding tissues with a plastic pestle on dry ice. The samples were sonicated for 15 seconds at 35 power level, then centrifugated at 14,000 rpm for 10 min at 4° C to remove undissolved substances. To denature the proteins, the lysates were boiled for 10 minutes with 1X Laemmli buffer, loaded into 4-20% gradient SDS-polyacrylamide gel electrophoresis, and the protein amounts were analyzed as described previously.^45^ We use primary antibody for RAB11B, β-catenin, and GAPDH incubated overnight in 3.5% BSA and secondary antibody conjugated to HRP for detecting protein levels.

### 4.6. Endosome quantification

We measured the number of apical endosomes in *Twist1* mutant and wild-type embryos to determine the impact of *Twist1* deletion on endocytic vesicle markers, LRP2 and RAB11B, and the number of endosomes. Immunofluorescent images of 3 different biological samples were collected for Wild type (WT), *Twist1* cko, and null embryos. Dual immunofluorescent staining for the endocytic factors LRP2 and β-catenin as well as RAB11B and β-catenin was performed to determine the location and number of the endocytic vesicles along the apical cell membrane. Using a multi-point tool in the software Image J^1^ version 5.3k, we manually counted the vesicles with red signals of 8 apical cells at either the dorsolateral hinge points and near midline in neural folds. A distinctly visible red fluorescent vesicle or yellow when overlaps with green β-catenin was counted as one vesicle, while for not so distinct red signal due to the densely packed endosomes, we avoided counting these vesicles unless we could estimate it two vesicles depending upon the size covered compared to a single vesicle.

### 4.7. Quantification of Cell-cell Junctions

Junction mapper software was used to quantify cell-cell junctions for apical cells at neural fold edges where pre-migratory CNCC usually emerged and delaminated.^48^ Cell-cell junctions were compared between WT and *Twist1* null and cko samples at E9.5 and E10.5. β-catenin was used as the junction marker for quantification due to prominent expression at the cell membrane. Interface contour, interface linearity index of neuroectodermal cells, interface area, and junction marker intensity per area were measured and compared with WT. Eight apical cells were picked within similar regions for comparison. Quantification was performed for cell-cell junctions of cells at the neural fold edges. Four cells directly along the apical membrane were chosen, along with the four cells directly behind them, for a total of 8 cells per region. Another method of choosing cells to quantify involved choosing eight cells directly along the apical membrane that were lined up in a linear manner. However, this method proved more difficult due to gaps and inconsistent spacing along the membrane, which led to more significant variation among the cells.

### 4.8. Statistical Analysis

All quantitative data are presented as the means ± standard deviations. Statistical comparisons were conducted using the unpaired student’s t-test for two groups, wild type and mutant, using GraphPad Prism 8 software. A significance level of p-value < 0.05 was considered statistically significant.

## Data Availability Statement

All data can be found within the article and its supporting information. Additional inquires can be directed to the authors of this study.

