## SUPPLEMENTARY FIGURES for "TWIST1 Modulates Cilia Length, Endocytic Vesicle Dynamics, and Cell-Cell Junctions during Neural Tube Morphogenesis"

**SUPPLEMENTARY FIGURE 1. Expression pattern of endocytic vesicle markers and adherens junction protein. (A-B''):** IF staining of LRP2 and β-catenin proteins in E9.5 *Twist1* null and WT littermate showing an increase in endosomal marker LRP2 in *Tw1 null* embryo compared to WT. **(C-D''):** IF staining of RAB11B and β-catenin proteins in E9.5 *Twist1* cko and WT littermate showing an increase in endosomal marker RAB11B in *Tw1 cko* embryo compared to WT.

**SUPPLEMENTARY FIGURE 2 Quantification analysis of cell-cell junction of neuroectodermal cells of *Twist1* null and WT embryos. (A):** IF staining of E9.5 WT mouse embryo using β-catenin as cell membrane marker. **(B):** IF staining of E9.5 *Twist1* null embryo using β-catenin marker. **(C, D):** Binary edge map of E9.5 WT and *Twist1* null embryos showing the cells used for quantification analysis. **(E-I):** Quantification analysis shows no significant change between *Twist1* null compared to WT due to large variations. **(J, K):** IF staining of E10 WT and *Twist1* null embryos using β-catenin. **(L, M):** Binary edge map of E10 WT and *Twist1* null mouse embryos showing the cells used for quantification analysis. **(N-R):** Quantification analysis shows a significant decrease in the interface contour, interface area, interface linearity index and JM1 intensity between *Twist1* null and WT embryos at E10.

**SUPPLEMENTARY FIGURE 3 Quantification analysis of cell-cell junction of neuroectodermal cells of Twist1 mutants and WT embryos. (A):** IF staining of E9.0 WT mouse embryo using β-catenin as cell membrane marker. **(B):** IF staining of E9.0 *Twist1* null embryo using β-catenin marker. **(C, D):** Binary edge map of E9.0 WT and *Twist1* null embryos showing the cells used for quantification analysis. **(E-I):** Quantification analysis shows no significant change between *Twist1* null compared to WT due to large variations. **(J, K):** IF staining of E9.5 WT and *Twist1* null embryos using β-catenin. **(L, M):** Binary edge map of E9.5 WT and *Twist1* null mouse embryos showing the cells used for quantification analysis. **(N-R):** Quantification analysis shows a significant increase in the interface linearity index between *Twist1* null and WT embryos at E9.5.

**SUPPLEMENTARY FIGURE 4 Quantification analysis of cell-cell junction of neuroectodermal cells of *Twist1* null and WT embryos. (A, B):** IF staining of E10 WT and *Twist1* null embryos using β-catenin. **(C, D):** Binary edge map of E10 WT and *Twist1* null mouse embryos showing the cells used for quantification analysis. **(E-I):** Quantification analysis shows a significant increase in the JM1 intensity area while there was no change in interface contour, interface linearity index, interface area, and JM1 intensity between *Twist1* null and WT embryos at E10.

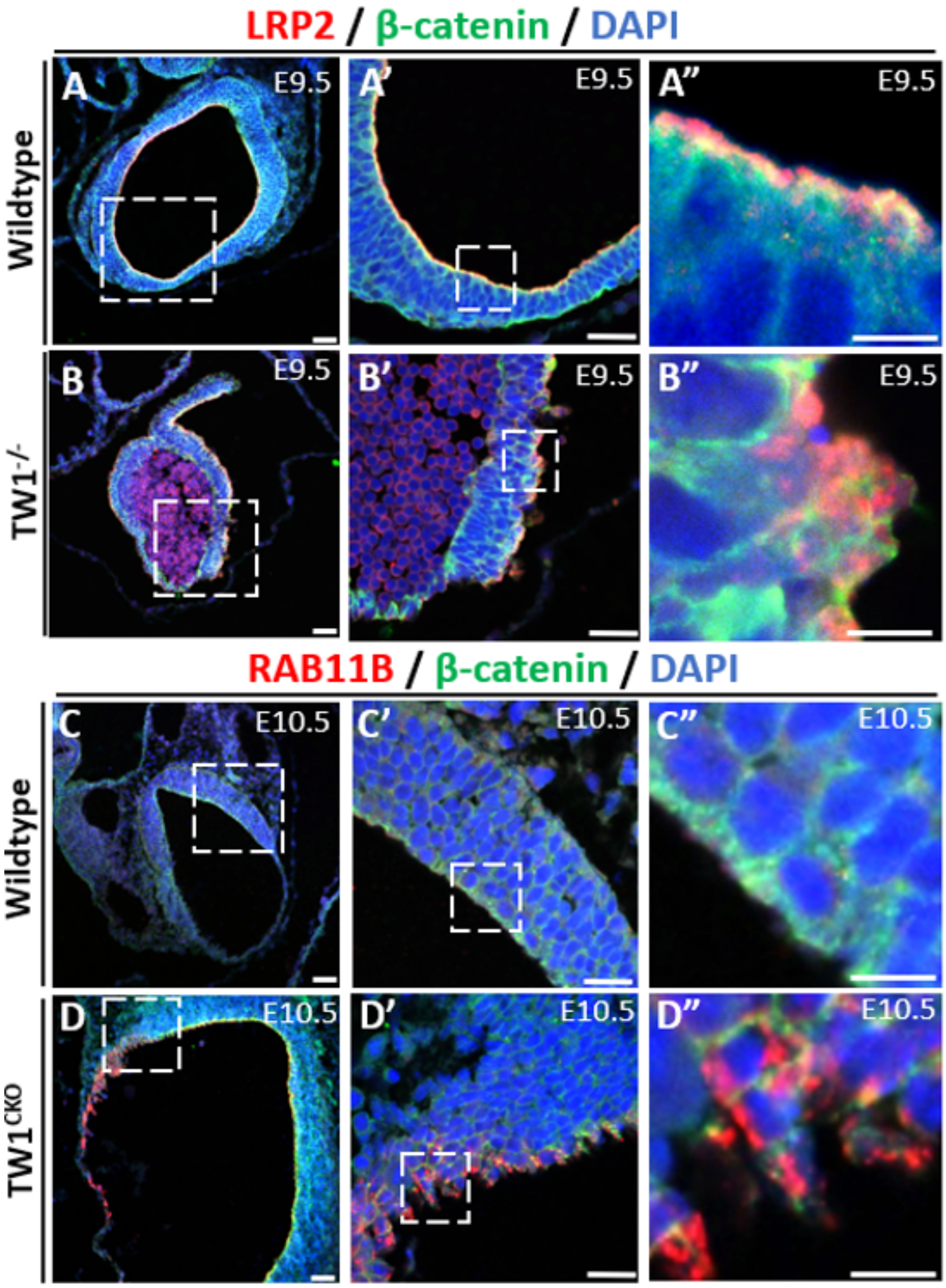


**SUPPLEMENTARY FIGURE 1.**

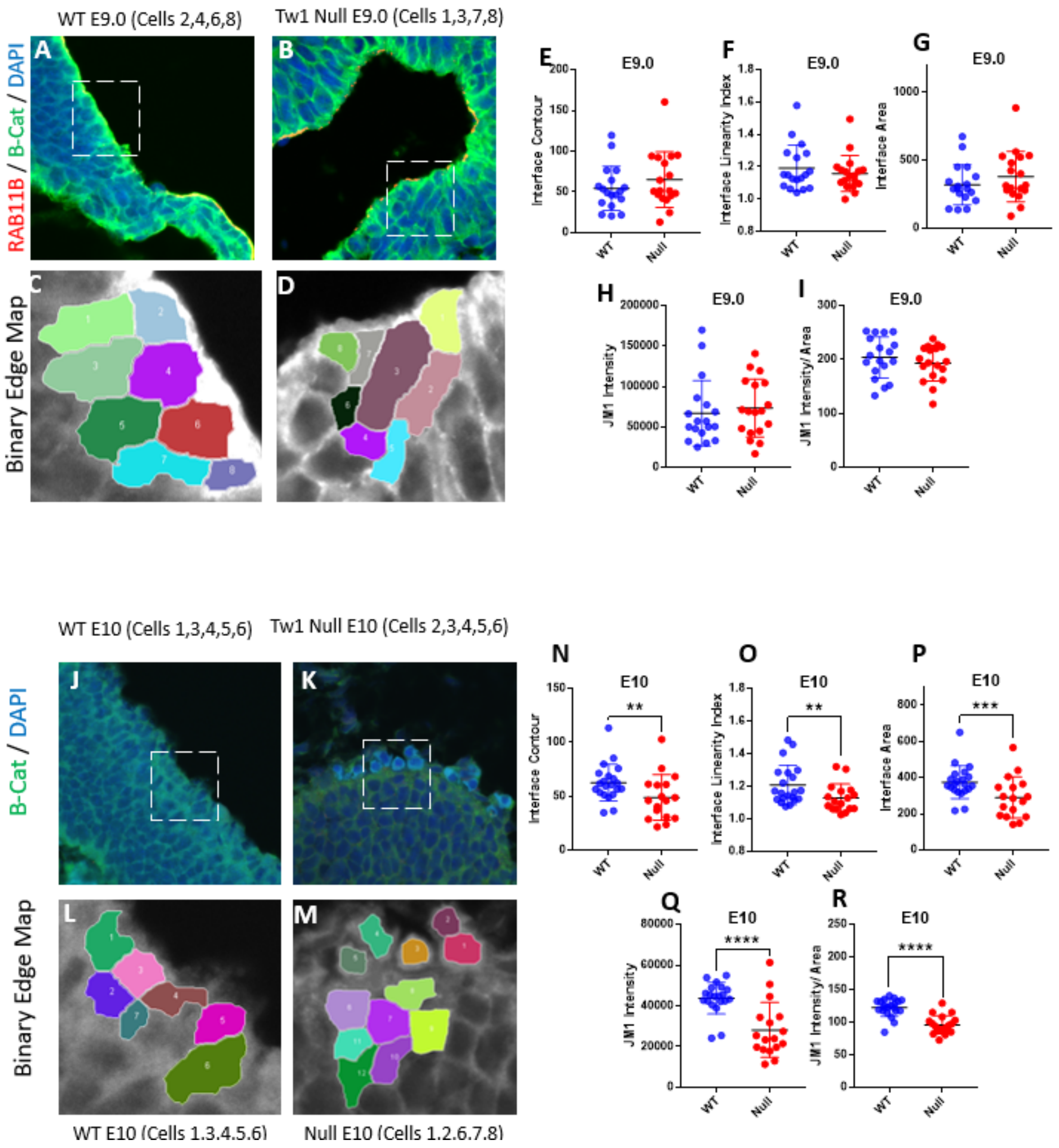


**SUPPLEMENTARY FIGURE 2.**

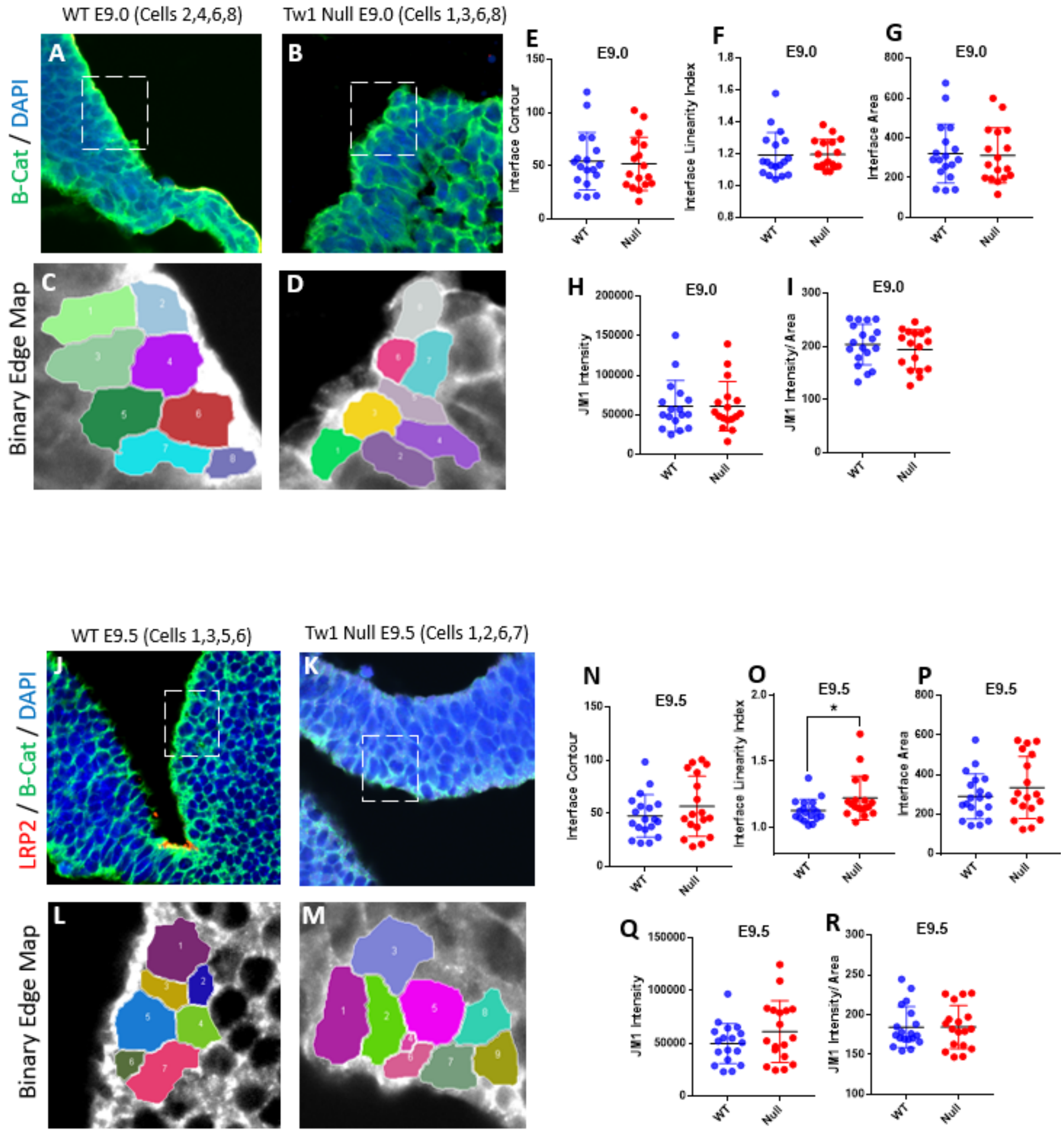


**SUPPLEMENTARY FIGURE 3**

.

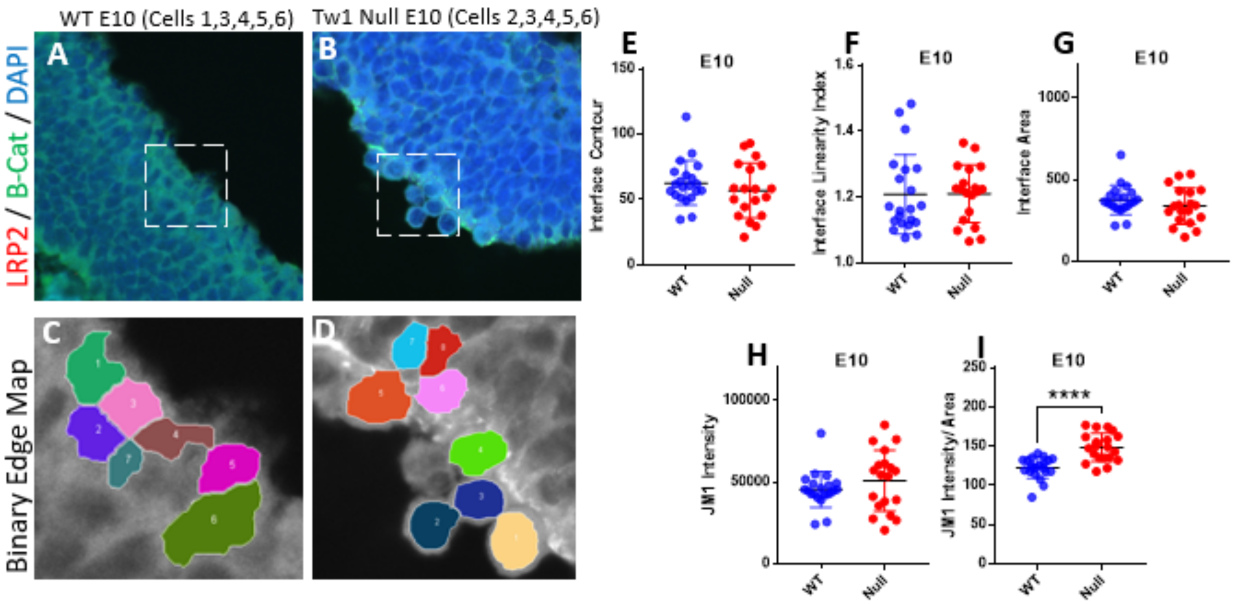


**SUPPLEMENTARY FIGURE 4.**
